# An integrated resource for systems-level analysis of aging hallmarks and associated genes

**DOI:** 10.64898/2026.05.29.728838

**Authors:** Rahul Tiwari, Mridhula Balaji, Nikhil Chivukula, Priyotosh Sil, Areejit Samal

**Author notes:** Corresponding author (A. Samal).

## Abstract

Aging is a complex biological process involving progressive cellular dysfunction, tissue decline, and increased susceptibility to multiple chronic diseases. A systemic view of aging through its established hallmarks provides a structured framework to understand this complexity and drive therapeutic discovery. To this end, we present AgingHallmarksDB, an interactive web platform that enables systems-level analysis of hallmark-associated gene sets. Aging-related genes were first curated from seven established resources, and those present in at least two of these resources were considered as consensus aging-related genes. Using functional annotations derived from GO, KEGG, and Reactome, a total of 3111 genes were mapped to the 11 aging hallmarks, of which 2593 were supported by additional experimental or manually curated evidence, with 1089 of these forming the consensus set. Further, AgingHallmarksDB supplements hallmarks and gene annotations with tissue or cell-type class specificity, exosomal profiles, and regulatory interactions, enabling users to perform hallmark enrichment, protein-protein and regulatory interaction-based analysis. To elucidate the interconnectedness of hallmarks, network separation and proximity analyses of hallmark-associated modules were performed, revealing functional overlap between hallmarks on the human interactome. Furthermore, the utility of AgingHallmarksDB was demonstrated through network topological and regulatory interaction analyses of common hallmark genes, hallmark enrichment and network proximity analyses of eight chronic age-related diseases and seven developmental and congenital diseases, and gene set enrichment analysis of a PM2.5-associated skin transcriptome. Together, these analyses highlight the utility of AgingHallmarksDB for aging hallmark-centred systems biology research. The resource is accessible at https://cb.imsc.res.in/aginghallmarksdb/.

## Introduction

Aging is characterized by the progressive accumulation of molecular damage and the breakdown of homeostasis, leading to a gradual decline in tissue and organismal function, reduced capacity to respond to stress [1], deterioration of both physical and mental integrity, ultimately increasing susceptibility to clinical complications and contributing to the natural progression toward the end of life [2]. Aging is one of the major demographic and biomedical challenges in the 21^st^ century. According to the World Health Organization (WHO), one in six people globally will be aged 60 years or older, with the number of individuals in this age group increasing from 1 billion in 2020 to 1.4 billion by 2030 [3]. By 2050, the global population aged 60 years and older is expected to increase to approximately 2.1 billion, while the number of people aged 80 years or older is projected to increase to 426 million [3]. This rapid demographic change is closely correlated with an increasing burden of age-related chronic diseases, including cardiovascular diseases, diabetes, cancer, neurological disorders, and musculoskeletal decline [3].

Although chronic diseases are strongly associated with aging, current research is progressively moving beyond a disease-oriented perspective, toward the broader objective of promoting healthy aging and longevity [4]. The WHO defines healthy aging as “the process of developing and maintaining the functional ability that enables well-being in older age” emphasizing that the focus of aging research can extend from disease management to the enhancement of healthspan and longevity [5]. In this context, frameworks describing the biological mechanisms of aging provide a valuable foundation for understanding how interconnected molecular and cellular processes collectively influence healthspan, longevity, and susceptibility to age-associated diseases.

Introduced by López-Otín and colleagues, the aging hallmarks framework conceptualizes the biological factors that collectively drive the aging phenotype. Initially proposed in 2013 [2] and later expanded in 2023 [6], the framework now encompasses twelve aging hallmarks which are: ‘Altered intercellular communication’, ‘Cellular senescence’, ‘Chronic inflammation’, ‘Deregulated nutrient-sensing’, ‘Disabled macroautophagy’, ‘Dysbiosis’, ‘Epigenetic alterations’, ‘Genomic instability’, ‘Loss of proteostasis’, ‘Mitochondrial dysfunction’, ‘Stem cell exhaustion’, and ‘Telomere attrition’. These hallmarks manifest naturally with age, speed up aging if they get worse, and improve health when fixed [2]. Since these hallmarks represent broad and interconnected biological processes rather than isolated entities, understanding their genetic complexity may provide deeper insights into aging-associated processes. Although Open Genes [7] curated aging hallmark- and longevity-associated genes, their integration of updated hallmark annotations and broader biological context remains limited. In particular, functional annotation and pathway-level information from resources such as KEGG [8] and Reactome [9] have not been integrated. Furthermore, gene-level annotations like cell or tissue presence, regulatory interactions and exosome presence remain dispersed across multiple resources. Moreover, hallmark-associated gene set enrichment analysis has highlighted intricate connections between genes and biological functions in the case of cancer hallmarks [10], while similar efforts are limited in the case of aging hallmarks.

To bridge this gap, we present AgingHallmarksDB, an integrative web platform that compiles and standardizes aging hallmark-associated genes, enabling a more systemic view of aging biology. The platform compiles aging associated-genes from seven databases, and functional annotations from Gene Ontology (GO), KEGG, and Reactome pathways to characterize biological functions associated with aging hallmarks. Additional levels of information, including cell-type classes or tissue-specific expression, exosome-associated expression, protein-protein, transcription factor (TF)-target, and kinase-substrate interactions, were integrated to provide broader biological context for hallmark-associated genes. The web platform enables users to perform aging hallmark enrichment analysis, interpret gene sets in the context of aging biology, and explore interconnected protein-protein and regulatory interactions, along with providing downloadable tables and networks for each. To understand interconnectedness or overlap of aging hallmarks at the level of the human interactome, network separation and proximity analyses were performed for hallmark-associated modules. Subsequently the practical use of AgingHallmarksDB, was shown through protein-protein interaction-based network topological and regulatory interaction analysis of common hallmark genes. Following that, the hallmark associations in aging-related chronic diseases and developmental and congenital diseases were shown through the over representation analysis (ORA)-based hallmark enrichment analysis, which was further supported through network proximity analysis. In addition, gene set enrichment analysis (GSEA) was performed using PM2.5-associated skin transcriptomic data. Together, these analyses provided systems-level insights into disease-aging and environment-aging contexts. Overall, AgingHallmarksDB is the first platform to provide a structured framework for studying aging through associated hallmarks, thereby supporting future longevity-focused research.

## Materials and Methods

### Curation of genes associated with aging hallmarks

We performed a systematic curation of human genes associated with the different aging hallmarks proposed by López-Otín *et al.* [2,6]. First, seven well-established aging-related databases, namely, Aging Atlas [11], AgingReG [12], CellAge [13], CSGene [14], Digital Ageing Atlas (DAA) [15], GenAge [13], and Open Genes [7] were selected based on their relevance and extensive coverage of genes implicated in aging processes. Gene-related information was retrieved and standardized based on Entrez identifiers using the mygene package (https://pypi.org/project/mygene/). These standardized gene sets were integrated to compile the ‘extended aging-related gene set’ comprising 5261 human genes, among which 1560 genes present in at least two of these seven databases were defined as the ‘consensus aging-related gene set’ (Supplementary Table S1).

López-Otín *et al.* [6] proposed twelve aging hallmarks that represented broad aging-related biological processes. To gain further insights, these hallmarks were manually annotated with 37 functional terms compiled from the analytical hallmarks used in Open Genes [7], and conceptual definitions provided by López-Otín *et al.* (Supplementary Table S2) [2,6], Subsequently, gene-associated functions, such as 160 GO terms classified as ‘Biological Process’ (BP) or ‘Molecular Function’ (MF) [16], 43 pathways from KEGG [8], and 65 pathways from Reactome [9], were manually mapped to the aging hallmarks based on the associated functional terms (Supplementary Table S2). Thereafter, the corresponding genes were mapped to the hallmarks through ‘gene2go.csv’ supplemented by QuickGO for GO term-based mapping, ‘NCBI2Reactome_All_Levels.csv’ for Reactome pathway-based mapping, KEGG API used for KEGG based mapping, along with the additional experimental or manual curation evidence obtained for GO and Reactome (Supplementary Table S3 and S4). In addition, genes associated with the sirtuin pathway were manually assigned to aging hallmark ‘Deregulated nutrient-sensing’ using HGNC annotations (https://www.genenames.org/) (Supplementary Table S3) [17]. Overall, the genes were assigned to the hallmarks with multilevel functional annotations. Applying the ≥2-database cutoff and filtering for experimentally validated and manually curated evidence resulted in four hallmark-associated gene datasets, extended and consensus gene sets, with and without evidence-based filtering (Supplementary Methods). Figure 1 summarizes the workflow followed in this study, while Table 1 summarizes the number of aging hallmark-associated genes curated from each resource.

**Figure 1.**
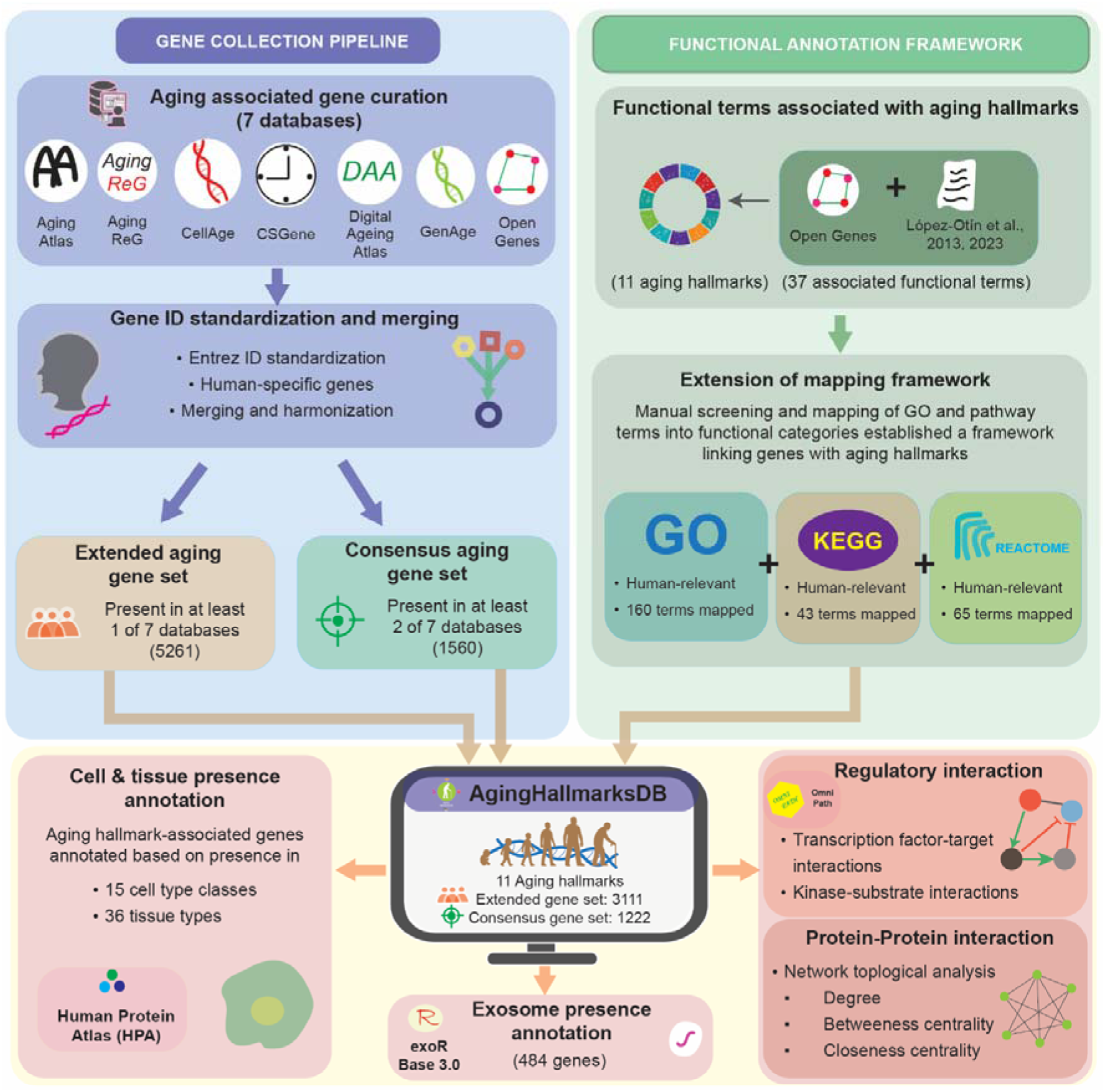
Schematic overview of the workflow used to develop AgingHallmarksDB.

**Table 1.** Number of aging-hallmark associated genes identified from GO, KEGG and Reactome. Note that KEGG does not provide a secondary evidence filter unlike other data sources, and therefore, pathway-gene mappings were utilized exactly as provided by KEGG.

| **Gene Set** | **Experimental or Manual Evidence of Gene-Function Association** | **Number of Genes Curated from GO** | **Number of Genes Curated from KEGG** | **Number of Genes Curated from Reactome** | **Number of Unique Genes** |
| --- | --- | --- | --- | --- | --- |
| Extended | Without Evidence Filter | 2403 | 1345 | 1551 | 3111 |
|  | With Evidence Filter | 1572 | 1345 | 1428 | 2593 |
| Consensus | Without Evidence Filter | 991 | 644 | 678 | 1222 |
|  | With Evidence Filter | 739 | 644 | 629 | 1089 |

### Additional characterization of hallmark-associated genes

To enable a systems-level understanding of hallmark-associated genes, additional information including single-cell expression, exosomal presence, and transcription factor (TF)-target and kinase-substrate interactions were retrieved. Single-cell expression data was obtained from the Human Protein Atlas [18] using ‘rna_single_cell_type.tsv’, and genes were mapped to 154 cell types based on an expression cut-off of nCPM > 1 (https://www.proteinatlas.org/humanproteome/single+cell/single+cell+type/method) [18]. The associated cell types were further categorized into 15 cell type classes and 36 tissues retrieved from HPA using ‘rna_single_cell_type_cell_types.tsv’ and ‘rna_single_cell_clusters.tsv’ resepectively (Supplementary Table S5). The exoRBase (v3.0) was utilized to identify the presence of hallmark-associated genes in exosomes; relevant annotations were sourced from ‘circRNAs_anno.csv’ and ‘longRNAs_anno.csv’ [19]. The TF-target and kinase-substrate phosphorylation interactions were sourced from DoRothEA and EnzSub datasets respectively of OmniPath [20], wherein the TF-target interactions were further filtered based on confidence scores A to C to obtain high-confidence interactions (Supplementary Methods).

### Curation of protein-protein interaction network

A large integrated human protein-protein interaction (PPI) network was curated by combining different comprehensive PPIs given by Gan *et al.*, Human interactome atlas and Ruiz *et al.* [21–23]. The resulting PPI network comprises 19572 nodes and 540192 edges (Supplementary Methods).

### Development of an integrated web platform

We created an integrated web platform, AgingHallmarksDB, which catalogs hallmark-associated genes. Further, AgingHallmarksDB enables hallmark enrichment analysis and provides network-based visualizations of TF-target and kinase-substrate interactions for user-supplied gene sets. The platform was developed using the Shiny web framework (https://shiny.posit.co/) in R (version 4.5.0) [24], and hosted on a webserver running on the Debian 13 Linux operating system. The user interface was developed using Bootstrap 5 (https://getbootstrap.com/docs/5.0/), while interactive data tables and network visualizations were implemented using DT (https://CRAN.R-project.org/package=DT) and visNetwork libraries (https://CRAN.R-project.org/package=visNetwork) in R. AgingHallmarksDB is openly accessible for research at https://cb.imsc.res.in/aginghallmarksdb/.

### Hallmark enrichment analysis tool

AgingHallmarksDB provides a ‘Hallmark Enrichment’ tool to enable users to identify which aging hallmarks are significantly represented in a user-supplied gene set using two different approaches, Over representation analysis (ORA) and Gene set enrichment analysis (GSEA) [25], Since the hallmark-associated genes can be both coding and non-coding, the gene sets annotated as ‘protein-coding’ and ‘ncRNA’ were obtained from the Bioconductor package org.Hs.eg.db [26], and utilized as the background gene set for the ORA. The statistical significance of gene overlap in ORA is assessed using a one-tailed hypergeometric test, implemented in R using the *phyper* function. To account for multiple testing, *p*-values are adjusted using the Benjamini-Hochberg (BH) correction [27]. Additionally, users can also perform GSEA-based enrichment analysis by providing gene expression fold-change values, enabling the identification of aging hallmarks and associated biological processes that are enriched among upregulated or downregulated genes [25]. Finally, the tool reports the overlapping gene sets, *p*-values, and adjusted *p*-values for each hallmark-associated gene set, and provides downloadable visualizations such as bar plots, bubble plots and circular plots.

### Curation of gene sets associated with chronic diseases and PM2.5 exposure

To showcase the utility of AgingHallmarksDB, we performed hallmark enrichment analysis for a panel of age-related chronic diseases, developmental and congenital diseases and PM2.5-exposed skin cells. Chronic diseases with known associations with aging-related biological processes were identified from Guo *et al* [28]. For each chronic age-related and developmental and congenital disease, the associated gene set was curated from DisGeNET using standardized disease names and relevant clinical subtypes (Supplementary Table S6) [29]. Subsequently, these associations were filtered to retain those with a gene-disease association score ≥ 0.4 (Supplementary Table S7). To ensure sufficient gene representation and reliable enrichment results, only diseases associated with at least 20 genes were retained for downstream analysis. The PM2.5 associated gene set was curated from an RNA-seq dataset (GSE143709) for PM2.5-treated normal human epidermal keratinocytes (NHEK) published by Kim *et al* [30].

### Assessment of the statistical significance of hallmark-associated largest connected component

The statistical significance of the largest connected component (LCC) on the curated human PPI network for each hallmark-associated gene set was evaluated using NetMedPy [31] (version 0.1.171) implemented in Python 3.11. For each hallmark, the observed LCC size was compared against a null distribution generated from 1000 randomly sampled gene sets of identical size and similar degree distribution using log-binning. The significance of the observed LCC was quantified using a z-score, where a larger positive z-score indicates that the hallmark-associated genes form a more connected module within the human interactome than expected by chance (Supplementary Methods).

### Network-based proximity and separation analysis

To investigate systems level interconnectedness or overlap of hallmark pairs, or hallmarks with age, developmental and congenital disease, the network-based analysis was performed using python package NetMedPy [31], on human PPI.

### Network separation analysis

For each pair of hallmark-associated module (LCC), PPI-based network separation analysis was performed using NetMedPy to quantify the topological relationships between different hallmark modules [31]. Lower (or negative) separation values indicate greater topological overlap or interconnectedness, whereas higher positive values indicate more distinct network localization (Supplementary Methods).

### Network proximity analysis

Quantifying the connections within the human interactome is necessary to comprehend the systems-level interaction between aging hallmarks and candidate gene sets. To this end, a large integrated human PPI network was subjected to network proximity analysis using NetMedPy [31]. Network proximity was quantified using the average minimum shortest path length. Symmetric proximity was calculated for hallmark-hallmark pairs, whereas asymmetric proximity was computed for hallmark-disease pairs. Statistical significance was assessed using degree-preserving randomization (log-binning) with 1000 iterations. A pair of gene sets is considered to be significantly close (or proximal), if the z-score was found to be negative with a *p*-value < 0.05 (Supplementary Methods).

## Results

### AgingHallmarksDB: A web platform for aging hallmark-associated genes

We present AgingHallmarksDB, an integrated web-based interactive platform that systematically integrates aging-associated genes from multiple aging-relevant databases and maps them to eleven of the twelve recognized aging hallmarks proposed by López-Otín *et al* [2,6]. Starting with age-associated genes compiled from seven distinct resources, an extended set of 5261 genes was obtained, of which 1560 genes present in at least two of seven databases, were defined as the consensus gene set in this study (Figure 1 and Supplementary Table S1) [7,11–15]. Functional annotation from GO (160 GO-Biological processes and Molecular function terms), KEGG (43 pathways), and Reactome (65 pathways) were used to systematically assign genes to hallmarks (Figure 1 and Supplementary Table S2 and S3). In total, 3111 genes were assigned to aging hallmarks, of which 1222 were present in the consensus gene set, thereby providing a structured resource for exploring molecular mechanisms underlying aging and age-related processes (Supplementary Table S3). Among these, 2593 genes were linked to aging hallmarks through KEGG pathways, GO terms and Reactome pathways that carried experimental or manually curated evidence for gene-to-term associations (Supplementary Table S4), of which 1089 were also present in the consensus aging-related gene set (Table 1 and Supplementary S8). This evidence-supported subset of 2593 genes were utilized as the primary hallmark-associated gene set for all downstream analyses in this study. For each hallmark-associated gene, AgingHallmarksDB provides several annotations, including associated cell type classes, tissues, and exosomal presence (Figure 1). AgingHallmarksDB is openly accessible for research at https://cb.imsc.res.in/aginghallmarksdb/..

### Distinct yet interconnected gene signatures across aging hallmarks

Comparison of hallmark-associated gene sets revealed that ‘Epigenetic alterations’ had the highest number (1198 genes) of associated genes, among which 545 genes were in the consensus set, whereas ‘Telomere attrition’ had the fewest (137 genes), among which 60 were present in the consensus set (Figures 2a and 2b). Furthermore, overlap analysis demonstrated interconnectedness among aging hallmarks, with ‘Epigenetic alterations’ and ‘Cellular senescence’ showing the highest overlap in the consensus set (201 shared genes), with the corresponding Jaccard index of 0.32, being the highest (Figures 2c and 2d). Furthermore, it was observed that ‘Epigenetic alterations’ contained the highest number (386) of unique hallmark-specific genes, of which 115 were in the consensus set, while ‘Telomere attrition’ contained the fewest (2) unique genes. (Supplementary Table S9). The complete overlap and Jaccard index across all hallmark-associated gene sets prior to evidence filtering is illustrated in Figure S1.

**Figure 2.**
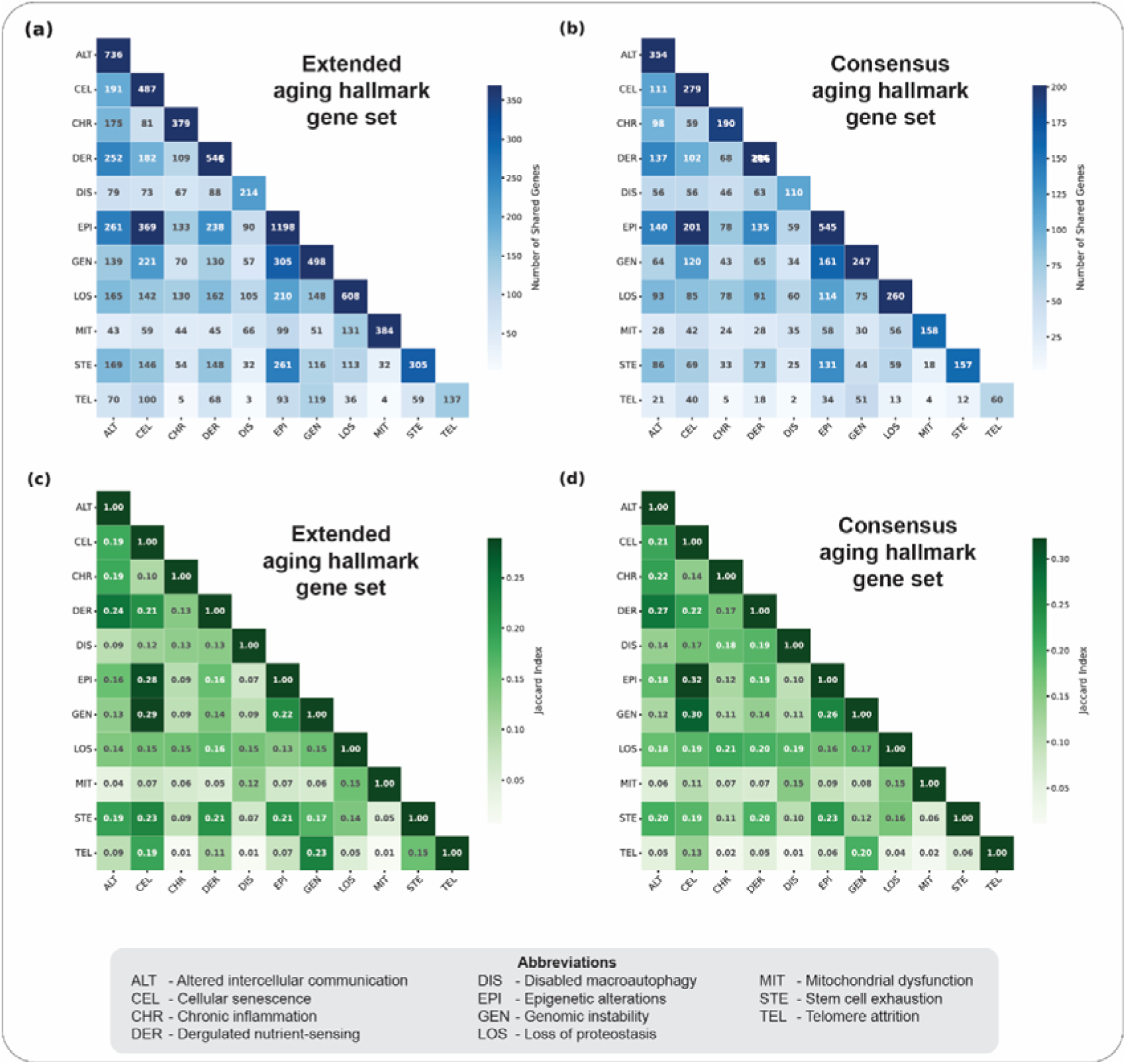
Pairwise analysis of gene overlaps between different aging hallmarks based on extended and consensus aging-related datasets comprising genes from KEGG pathways and those with experimental or manual evidence of association from GO and Reactome. **(a)** Pairwise overlap between aging hallmark-associated extended gene sets. **(b)** Pairwise overlap between aging hallmark-associated consensus gene sets. **(c)** Pairwise Jaccard index between aging hallmark-associated extended gene sets. **(d)** Pairwise Jaccard index between aging hallmark-associated consensus gene sets.

### Aging hallmark-associated genes show broad distribution across diverse cell type classes and tissues

Among 15 cell-type classes, specialized epithelial cells showed the highest association with 2454 genes, of which 1043 were present in the consensus set, whereas trophoblast cells showed the lowest with 2070 genes, of which 909 were in the consensus set (Supplementary Table S10). Among the 36 tissue types associated with aging related genes, testis had the highest number (2510), of which 1060 were in the consensus set, while brain neuron had the least (2270), of which 964 were in the consensus set (Supplementary Table S10). These results highlight the broad cellular and tissue-specific distribution of aging hallmark-associated genes.

### Exosomal annotation of aging-associated genes highlights their extracellular relevance in aging

Mapping of hallmark-associated genes to exosome-related datasets revealed 397 exosome-associated genes of which 181 genes were present in the consensus set. Among the hallmarks, ‘Epigenetic alterations’ showed the highest number (178) of exosome-associated genes, of which 91 genes were in the consensus set (Supplementary Table S8), highlighting the potential role of exosomal cargo in epigenetic alterations associated with aging [32].

### Regulatory interactions reveal transcriptional and post-translational control across aging hallmarks

Transcription factor (TF)-target and kinase-substrate annotations were integrated into the hallmark-associated gene sets to characterize their regulatory interactions. Among the 2593 aging hallmark-associated genes, 168 were TFs that were connected to 1193 target genes, while 557 were kinases that were connected to 1116 substrate proteins (Supplementary Table S11). Within the 1089 genes in the consensus set, 100 were TFs, connected to 574 target genes and 326 were kinases connected to 571 substrate proteins (Supplementary Table S11). Together, these observations indicate the diverse transcriptional and post-translational regulatory interactions underlying aging hallmarks.

### AgingHallmarksDB web interface enables interactive hallmark-based gene exploration

AgingHallmarksDB provides an interactive web interface for exploring aging hallmark-associated genes and their functional characteristics (Figures 3 and 4).

**Figure 3.**
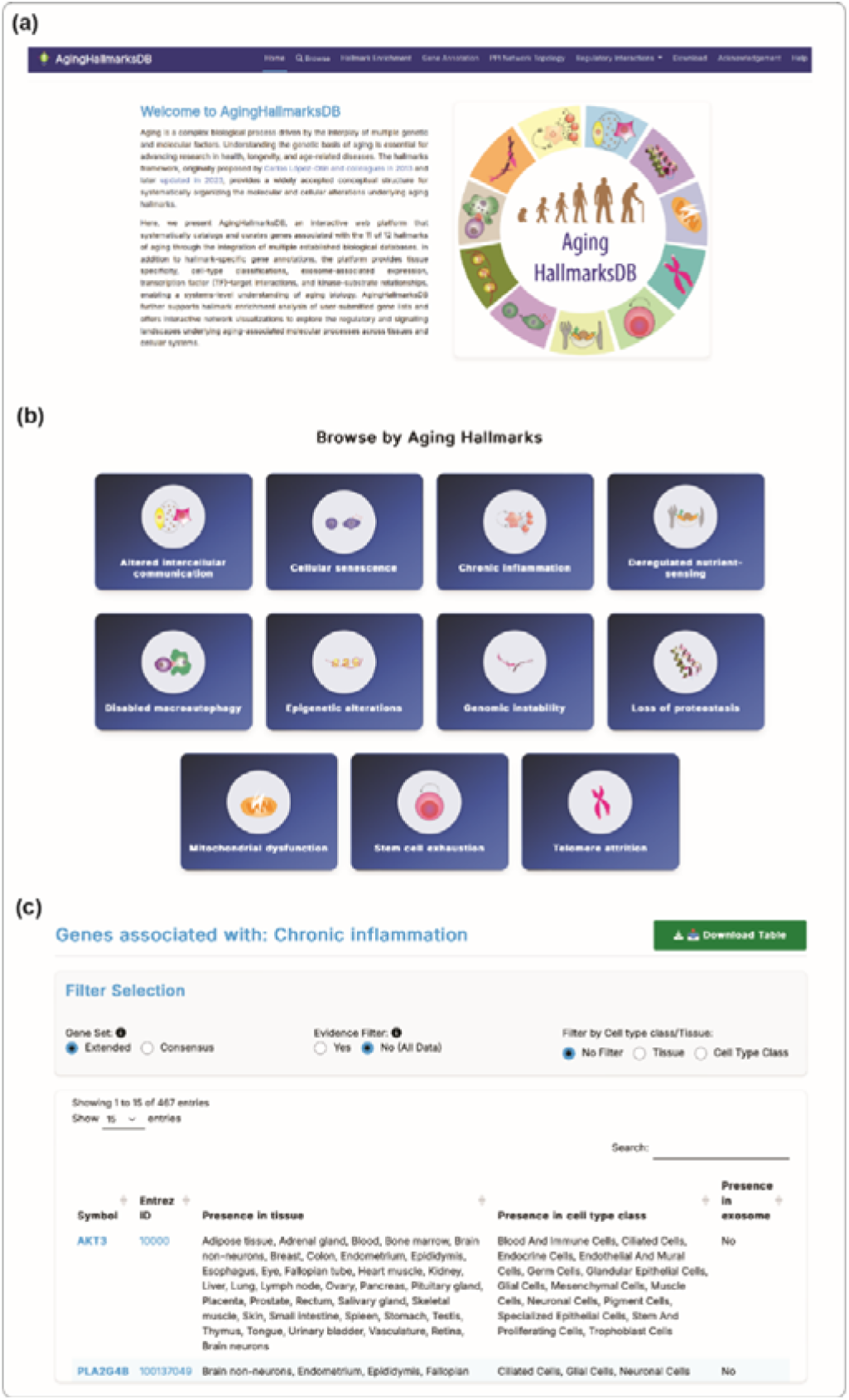
Web interface of AgingHallmarksDB. **(a)** ‘Home’ section, which provides an overview of the platform and summarizes its major features and functional modules. **(b)** ‘Browse’ section, where users can explore genes associated with the 11 aging hallmarks. **(c)** Gene information page associated with ‘Chronic inflammation’.

**Figure 4.**
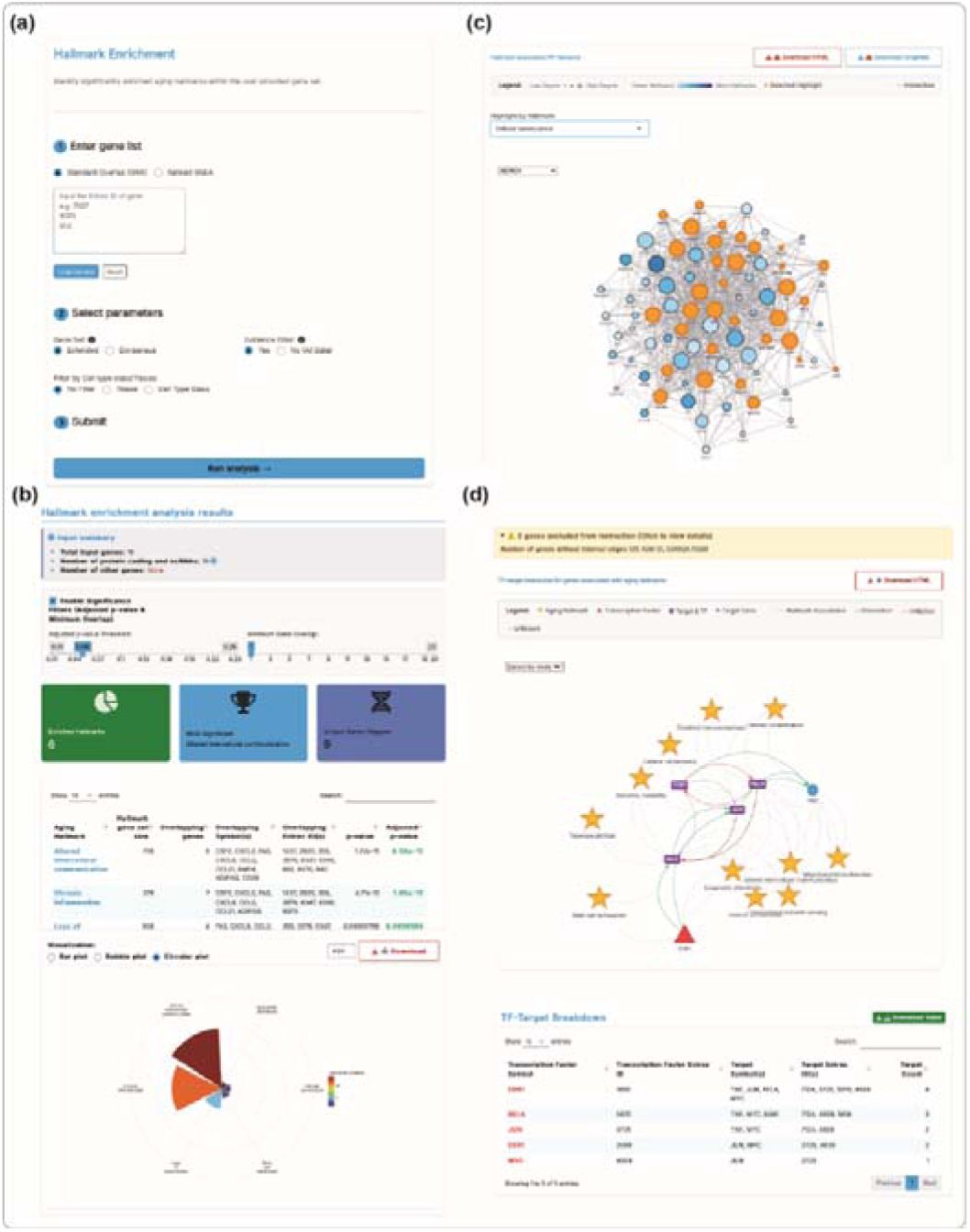
Hallmark enrichment and network visualization tools in AgingHallmarksDB. **(a)** Input page for hallmark enrichment analysis with options to select gene set, evidence filter, tissue and cell type class filtering. **(b)** Results page of ORA-based enrichment analysis showing input summary, significance filters, enriched hallmarks, most significant hallmark and downloadable result tables. **(c)** Protein-Protein interaction network visualization showing the interactions between different hallmark associated genes for an input gene set. The interactive network can be exported along with node and edge lists. **(d)** Regulatory network visualization showing transcription factor (TF)-target interactions connected to aging hallmark-associated genes, the interactive figure in downloadable format, followed by the TF-target results table.

### Browse

The ‘Browse’ section allows users to navigate individual aging hallmarks and access their associated genes together with tissue and exosome annotations (Figure 3b). The interface supports interactive exploration of hallmark-specific datasets and provides downloadable CSV files for downstream analysis (Figure 3c).

### Hallmark enrichment

The ‘Hallmark Enrichment’ section allows users to submit a custom gene set to identify significantly associated aging hallmarks (Figure 4a). This section comprises two enrichment analysis options: the hypergeometric test-based over-representation analysis (ORA) and gene set enrichment analysis (GSEA), allowing users to select the approach most appropriate for their data and research objectives. The analysis generates a downloadable enrichment table containing detailed statistical information and provides multiple visualization options, including bar plots, bubble plots, and circular plots (Figure 4b). Users can further customize the analysis by applying adjustable cutoffs for adjusted *p*-values and gene overlap to prioritize the most relevant aging hallmarks associated with their input gene set.

### Gene annotation

The ‘Gene Annotation’ section enables users to annotate custom gene sets with cell-type class, tissue-specific expression, exosome-associated information, and associated aging hallmarks. This feature helps users explore the cellular and biological context of their genes of interest through integrated annotation datasets.

### PPI network topology

The ‘PPI Network Topology’ section enables users to upload a gene set and explore its corresponding protein-protein interaction (PPI) network. The platform provides network topology-based insights by calculating key node properties, including degree, betweenness centrality, and closeness centrality. The tab also provides downloadable networks in the form of edge and node lists, along with GraphML format (Figure 4c).

### Regulatory interactions

The ‘Regulatory Interactions’ section allows users to upload a gene set and explore its TF-target and kinase-substrate interactions. The platform provides interactive regulatory network visualization along with downloadable annotation results (Figure 4d), enabling users to investigate transcriptional and post-translational regulatory relationships associated with their genes of interest.

### Comparison of AgingHallmarksDB with an existing hallmarks resource

Open Genes[7] provides aging-associated genes and includes 23 analytical hallmarks mainly annotated using GO Biological Process terms, covering 1224 genes associated to hallmarks of aging. In comparison, AgingHallmarksDB offers a broader hallmark-centered framework with 37 hallmark-associated functional terms annotated using GO Biological Process and Molecular Function terms, along with KEGG and Reactome pathways. The database expands the coverage to 3111 hallmark-associated genes and additionally incorporates non-coding RNAs with user-selectable evidence filtering options. Further, AgingHallmarksDB extends beyond conventional hallmark annotation by integrating tissue and cell-type information, exosome-associated expression profiles, TF-target interactions, and kinase-substrate relationships, thereby enabling systems-level exploration of aging biology. In addition, the platform includes a dedicated hallmark enrichment tool that identifies significantly associated aging hallmarks for user-supplied gene sets, provides visualization through enrichment plots and enables exploration of regulatory interactions. It was observed that 126 genes that were previously associated with aging hallmarks in Open Genes, could not be mapped to any hallmark in AgingHallmarksDB. This can be attributed to the substantial changes in the GO annotations used for hallmark assignment, which not only introduced new gene associations, but also removed some pre-existing terms. The full comparison table between Open Genes and AgingHallmarksDB is provided here as Table 2.

**Table 2.**
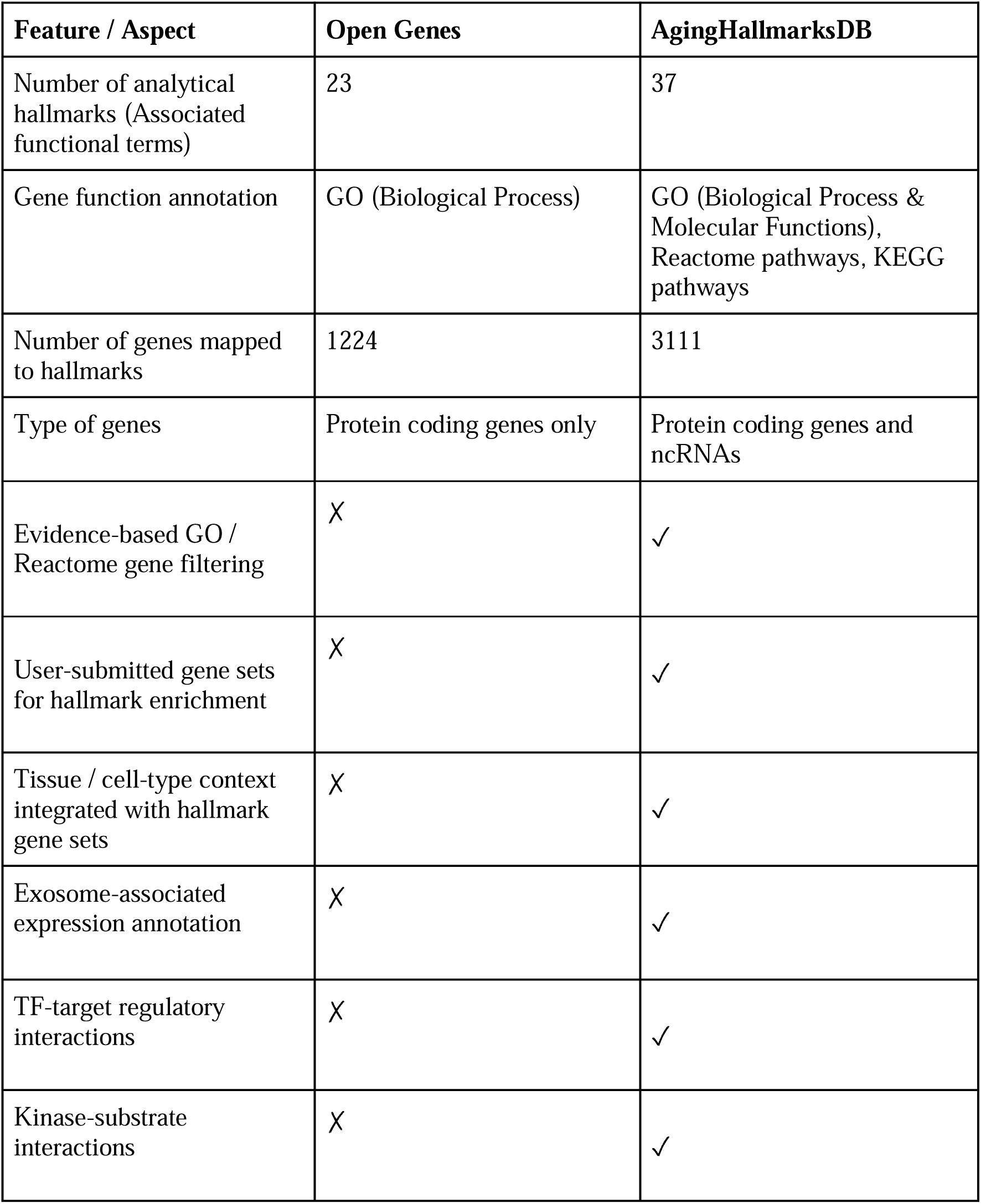
Comparative overview of Open Genes and AgingHallmarksDB.

### Aging hallmark-associated modules display high topological overlap in the human protein-protein interaction network

Aging hallmarks represent biologically interconnected processes that collectively drive aging phenotypes. To investigate their systems-level connectivity, hallmark-associated modules were constructed by extracting the largest connected component (LCC) of the consensus aging hallmark gene sets (with evidence), followed by network separation and network proximity analyses. Significant modules were successfully identified for all 11 aging hallmarks (z-score > 1.96, *p*-value < 0.05), with module significance z-scores ranging from 2.32 to 3.98 (Supplementary Figure S2). The ‘Telomere attrition’ module, comprising 59 genes, exhibited the highest significance (z-score = 3.98), whereas the ‘Loss of proteostasis’ module showed the lowest significance (z-score = 2.32) (Supplementary Figure S2; Table S12).

Network separation analysis was subsequently performed to assess the topological overlap among hallmark-associated modules within the PPI network. The analysis demonstrated that 10 of the 11 hallmark modules substantially overlapped with one another, as indicated by negative separation values. The strongest overlap was observed between ‘Epigenetic alterations’ and ‘Cellular senescence’ (separation = -0.48), followed by ‘Genomic instability’ and ‘Cellular senescence’ (separation = -0.45). In contrast, the ‘Telomere attrition’ module overlapped only with ‘Cellular senescence’ (separation = -0.11) and ‘Genomic instability’ (separation = -0.27) (Figure 5a).

**Figure 5.**
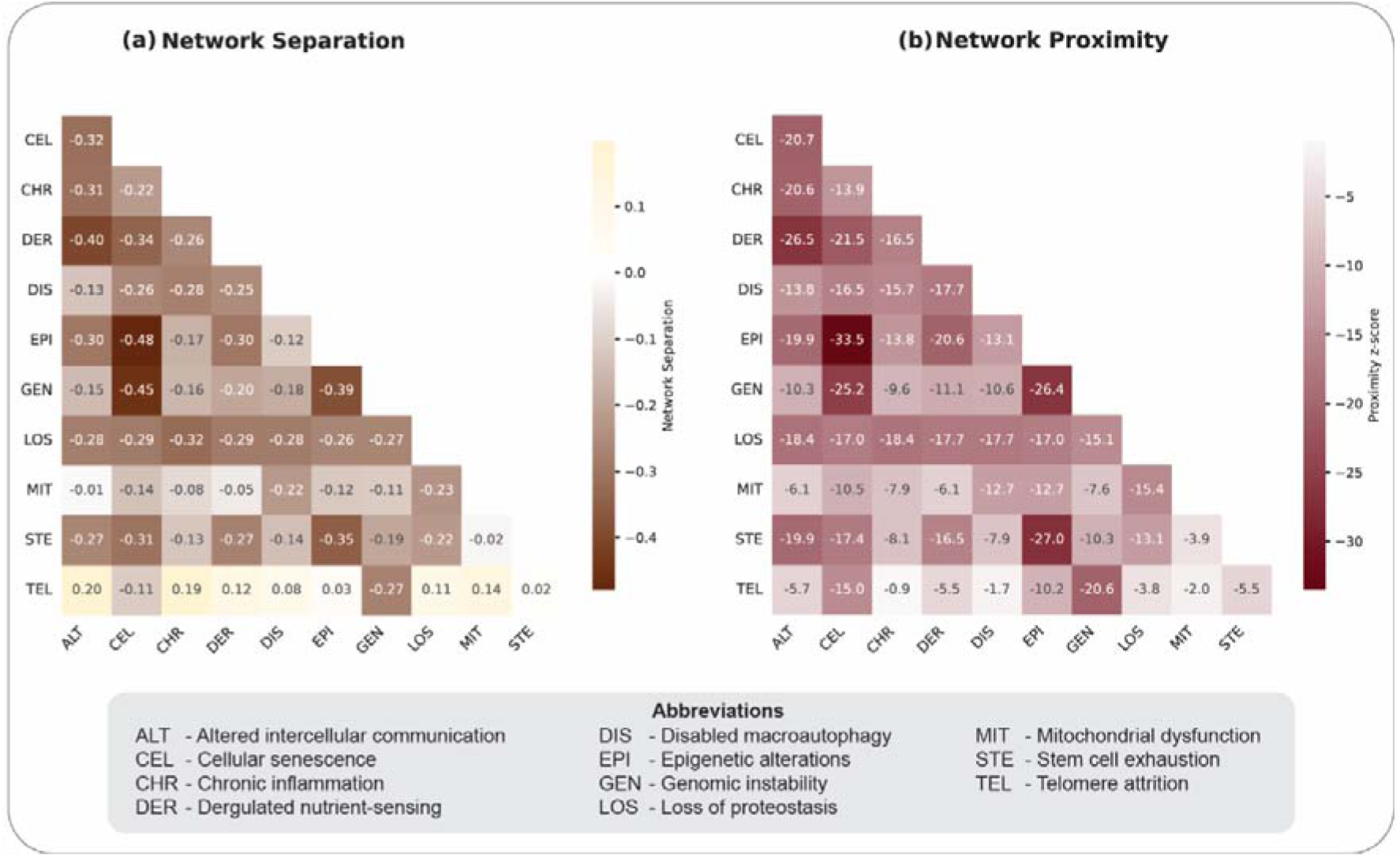
Human interactome based separation and proximity analysis of overlaps between different pairs of aging hallmark modules for consensus aging hallmark-associated datasets. **(a)** Pairwise separation between aging hallmark-associated modules. **(b)** Pairwise proximity between aging hallmark-associated modules.

Consistent with the network separation results, complementary network proximity analysis further demonstrated extensive network proximity among hallmark modules. ‘Epigenetic alterations’ and ‘Cellular senescence’ exhibited the strongest proximity (z-score = -33.5), followed by ‘Stem cell exhaustion’ and ‘Epigenetic alterations’ (z-score = -27.0), ‘Genomic instability’ and ‘Epigenetic alterations’ (z-score = -26.4), and ‘Genomic instability’ and ‘Cellular senescence’ (z-score = -25.2). Almost every hallmark pair exhibited significant network proximity (Figure 5b).

### Common genes reveal the interconnectedness of aging hallmarks

Aging hallmarks are biologically interconnected, and this interconnection is, in part, driven by genes shared across multiple hallmarks. Analysis identified 22 genes that were common to at least eight of the 11 aging hallmark modules for consensus aging hallmark associated gene set (with evidence), suggesting that these genes may collectively contribute to multiple hallmark processes (Supplementary Table S13). To further characterize their biological significance, PPI-based topological analysis was performed using the ‘PPI Network Topology’ section of AgingHallmarksDB. Among the shared genes, UBC emerged as the highest-ranked node based on degree (20), betweenness centrality (0.3892), and closeness centrality (0.9545), followed by TP53, MAPK1, GSK3B, and AKT1 (Figure 6a), indicating their central roles within the aging interaction network (Supplementary Table S13). Notably, multiple members of the MAPK family, including MAPK1, MAPK3, and MAPK8 (Figure 6a), were also among the shared multi-hallmark genes. Subsequently, the regulatory potential of these common hallmark genes was investigated using the ‘Regulatory Interaction’ section of AgingHallmarksDB. This analysis identified TP53 as both a transcription factor and a phosphorylation regulatory node (Figure 6b,c). Additionally, RELA and NFKB1 were identified as key regulatory nodes within the transcription factor-target interaction network (Figure 6b,c).

**Figure 6.**
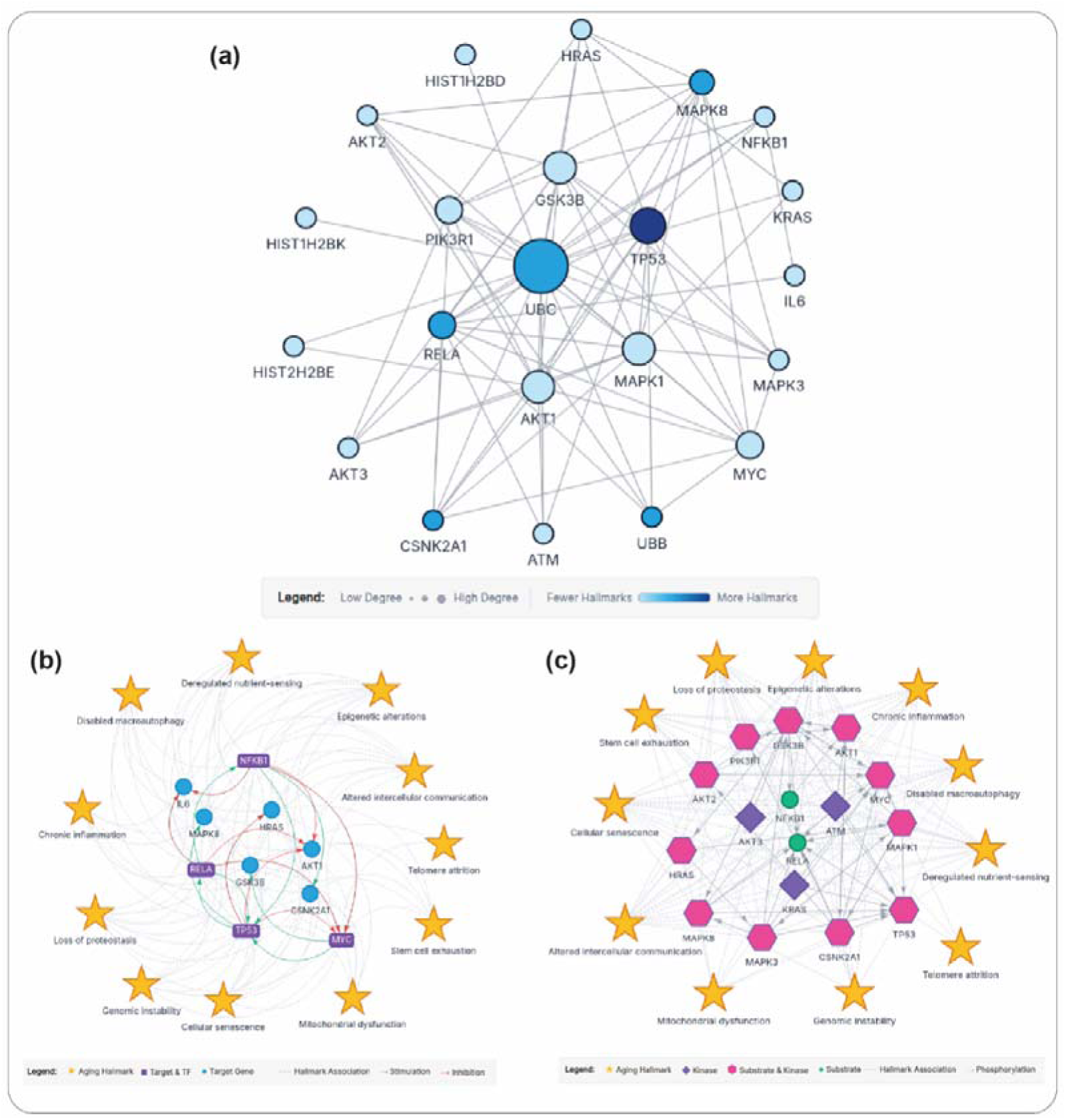
Human protein-protein interaction (PPI), transcription factor-target, and kinase-substrate interaction networks of common genes associated with at least eight of the 11 aging hallmarks. **(a)** PPI network of the common hallmark genes. Node size is proportional to degree, and node color intensity indicates the number of aging hallmarks with which each gene is associated. **(b)** Transcription factor-target interaction network of the common hallmark genes. Violet nodes represent genes that function as both transcription factors and target genes, whereas blue nodes represent target genes only. **(c)** Kinase-substrate interaction network of the common hallmark genes. Pink nodes represent genes that function as both kinases and substrates, whereas violet nodes represent genes that function only as kinases.

### Aging hallmarks enrichment patterns distinguish chronic age-associated diseases from developmental and congenital diseases

Aging hallmarks represent interconnected biological processes that describe the onset and progression of aging. While developmental and congenital diseases may share certain molecular and biological pathways with aging-related processes, they primarily originate from perturbations in developmental processes rather than age-associated biological decline. Therefore, comparison of enrichment patterns between these two groups enabled assessment of whether AgingHallmarksDB preferentially captures hallmark-level signatures associated with aging-related disease biology. To investigate this relationship, gene sets for chronic age-associated diseases and developmental and congenital diseases were curated from DisGeNET (Supplementary Table S6 and S7). Age-associated diseases, including Alzheimer’s disease, Parkinson’s disease, heart failure, COPD, and type 2 diabetes mellitus, showed extensive enrichment across the aging hallmarks, with all diseases exhibiting enrichment for 10 or 11 of the 11 hallmarks. In contrast, developmental and congenital diseases displayed substantially lower levels of enrichment, typically involving only 0-4 hallmarks (Table 3 and Supplementary Table S14). For example, polydactyly showed no significant hallmark enrichment, while epidermolysis bullosa and spinal muscular atrophy were enriched for only a single hallmark (Supplementary Table S14). These findings indicate that AgingHallmarksDB captures biological processes associated with aging and age-related diseases rather than broadly enriching disease-associated gene sets, thereby demonstrating its ability to distinguish aging-associated conditions from diseases primarily arising from developmental and genetic perturbations.

**Table 3.**
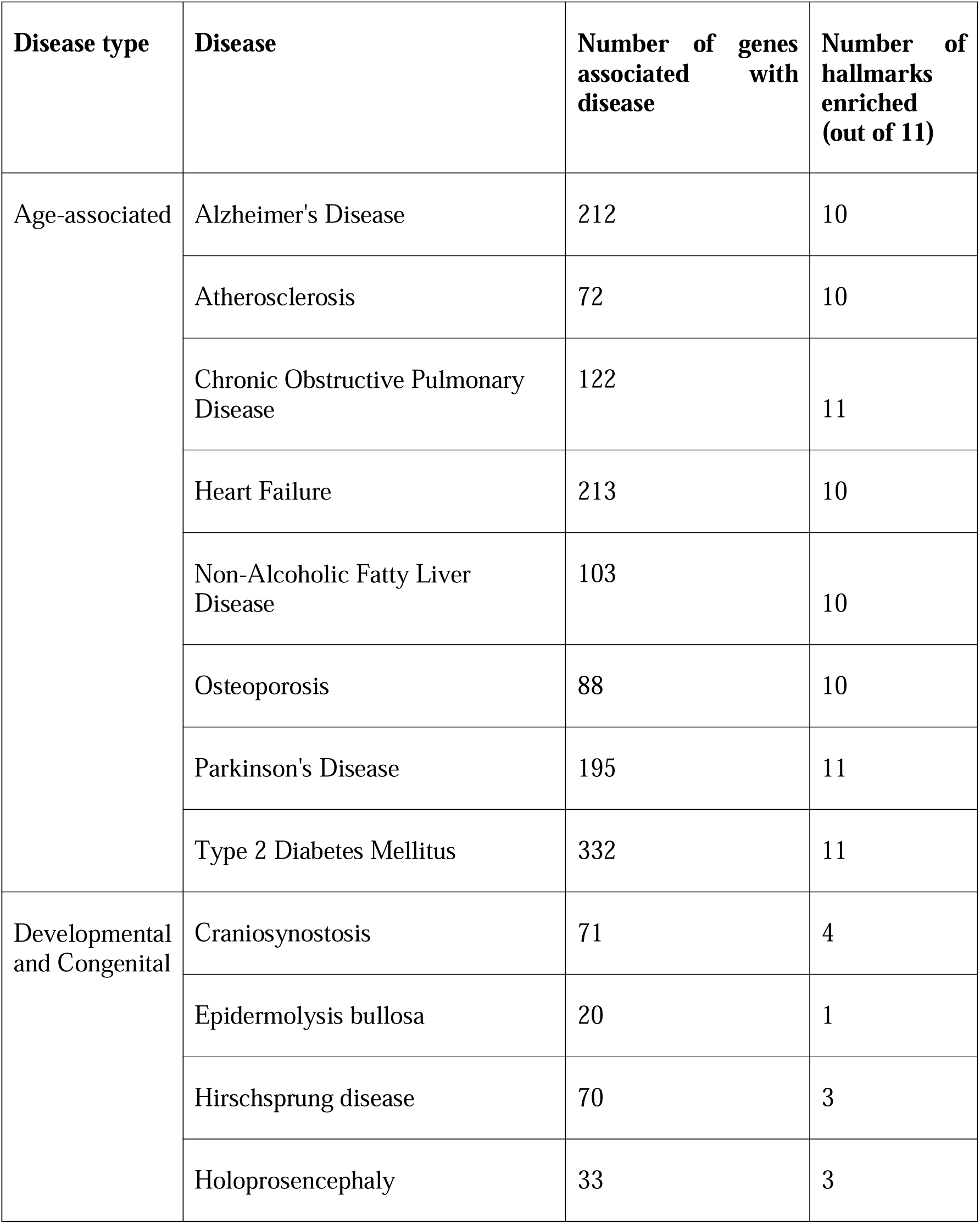
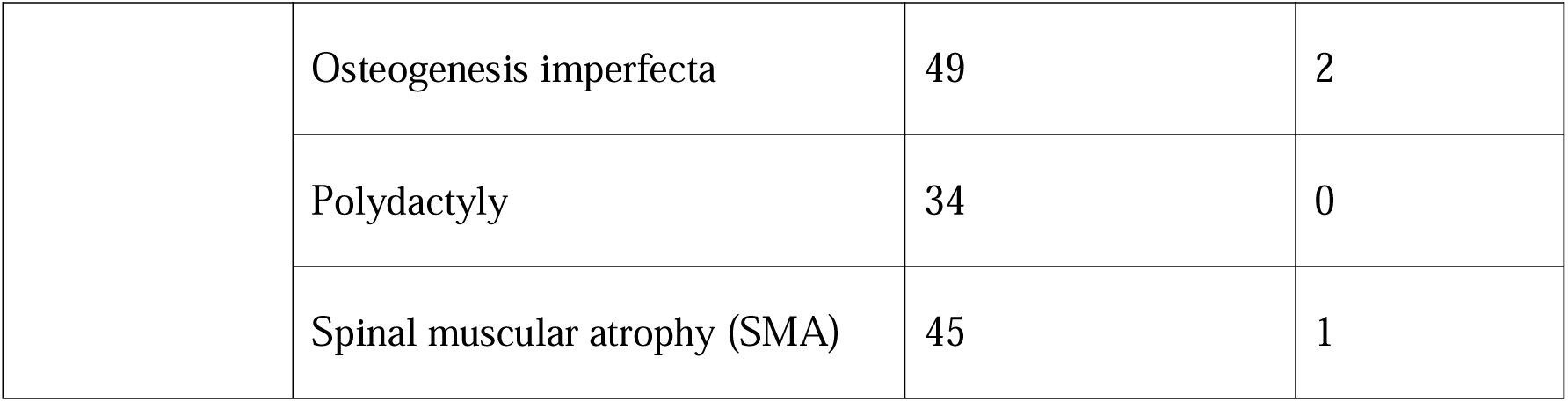
Comparative aging hallmark enrichment in age-associated and developmental and congenital disease gene sets.

| **Disease type** | **Disease** | **Number of genes associated with disease** | **Number of hallmarks enriched (out of 11)** |
| --- | --- | --- | --- |
| Age-associated | Alzheimer's Disease | 212 | 10 |
|  | Atherosclerosis | 72 | 10 |
|  | Chronic Obstructive Pulmonary Disease | 122 | 11 |
|  | Heart Failure | 213 | 10 |
|  | Non-Alcoholic Fatty Liver Disease | 103 | 10 |
|  | Osteoporosis | 88 | 10 |
|  | Parkinson's Disease | 195 | 11 |
|  | Type 2 Diabetes Mellitus | 332 | 11 |
| Developmental and Congenital | Craniosynostosis | 71 | 4 |
|  | Epidermolysis bullosa | 20 | 1 |
|  | Hirschsprung disease | 70 | 3 |
|  | Holoprosencephaly | 33 | 3 |
|  | Osteogenesis imperfecta | 49 | 2 |
|  | Polydactyly | 34 | 0 |
|  | Spinal muscular atrophy (SMA) | 45 | 1 |

### Hallmark signature profiling using over representation analysis reveals associations between aging and chronic disease progression

The strong association of aging-associated diseases with aging hallmarks provides a systems-level view of chronic diseases. For example, in Alzheimer’s disease, ‘Loss of proteostasis’ was the most significantly enriched hallmark (adjusted *p*-value = 1.53E-35) (Figure 7a), which was followed by ‘Deregulated nutrient-sensing’ (adjusted *p*-value = 2.13E-24) (Figure 7a and Supplementary Table S14). Network proximity analysis further revealed that ‘Chronic inflammation’ has the strongest association with Alzheimer’s disease (z-score = −6.54, p-value < 0.001), Consistent with the enrichment analysis, ‘Loss of proteostasis’ also demonstrated a strong association with Alzheimer’s disease in network proximity analysis as well, which in turn further supports its potential role in disease pathogenesis further supporting its potential role in disease pathogenesis (z-score = −6.36, p-value < 0.001) (Figure 7b and Supplementary Table S15).

**Figure 7.**
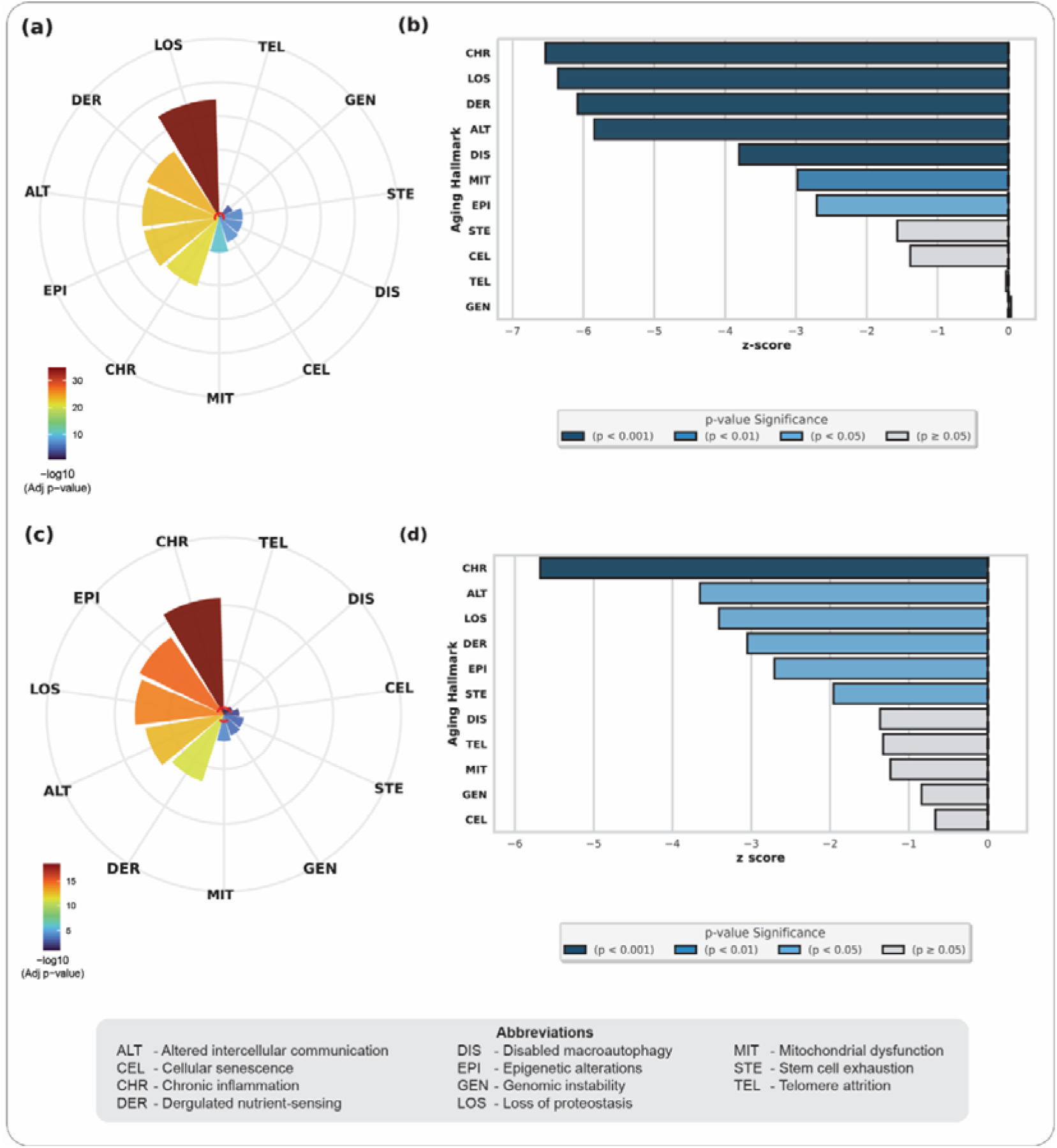
Hallmark enrichment and network proximity analysis results for Alzheimer’s disease and atherosclerosis. **(a)** Circular enrichment plot for Alzheimer’s disease. **(b)** Network proximity bar plot for Alzheimer’s disease-associated genes against aging hallmark-associated consensus gene sets. **(c)** Circular enrichment plot for atherosclerosis. **(d)** Network proximity bar plot for atherosclerosis-associated genes against aging hallmark-associated consensus gene sets.

Similarly, in Atherosclerosis, ‘Chronic inflammation’ emerged as the most significantly enriched hallmark (adjusted *p*-value = 4.29E−22) (Figure 7c and Supplementary Table S14). Network proximity-based analysis also identified ‘Chronic inflammation’ as the closest hallmark to the atherosclerosis-associated gene set (z-score = −5.68, *p*-value < 0.001) (Figure 7d and Supplementary Table S15). The remaining diseases also exhibited distinct hallmark signature profiles, highlighting the predominant aging-associated biological processes linked to each condition (Supplementary Tables S14 and S15). Collectively, these observations support the biological relevance of hallmark-based enrichment analysis in characterizing aging-associated disease mechanisms.

### Skin tissue-based enrichment of PM2.5-associated genes revealed epigenetic alterations as a key contributor

To investigate the relationship between exposure to fine particulate matter with aging hallmarks, PM2.5-associated genes were analyzed using AgingHallmarksDB in the context of skin aging. Gene set enrichment analysis (GSEA) of PM2.5 exposure-associated genes against the aging hallmark gene set (consensus with evidence for skin tissue) revealed significant enrichment of 10 of 11 skin-related aging hallmarks. Among these, ‘Epigenetic alterations’ exhibited the strongest enrichment signal (NES = −2.901, adjusted p-value = 1.52E-7), followed by ‘Chronic inflammation’ (NES = 3.47, adjusted p-value = 5.977E-5) (Supplementary Table S16). The negative and positive enrichment scores reflect the direction of enrichment of genes within the ranked PM2.5-associated gene list. Together, these findings highlight the enrichment of epigenetic alterations- and chronic inflammation-associated genes among the skin-related aging hallmarks identified in the PM2.5 exposure signature (Figure 8).

**Figure 8.**
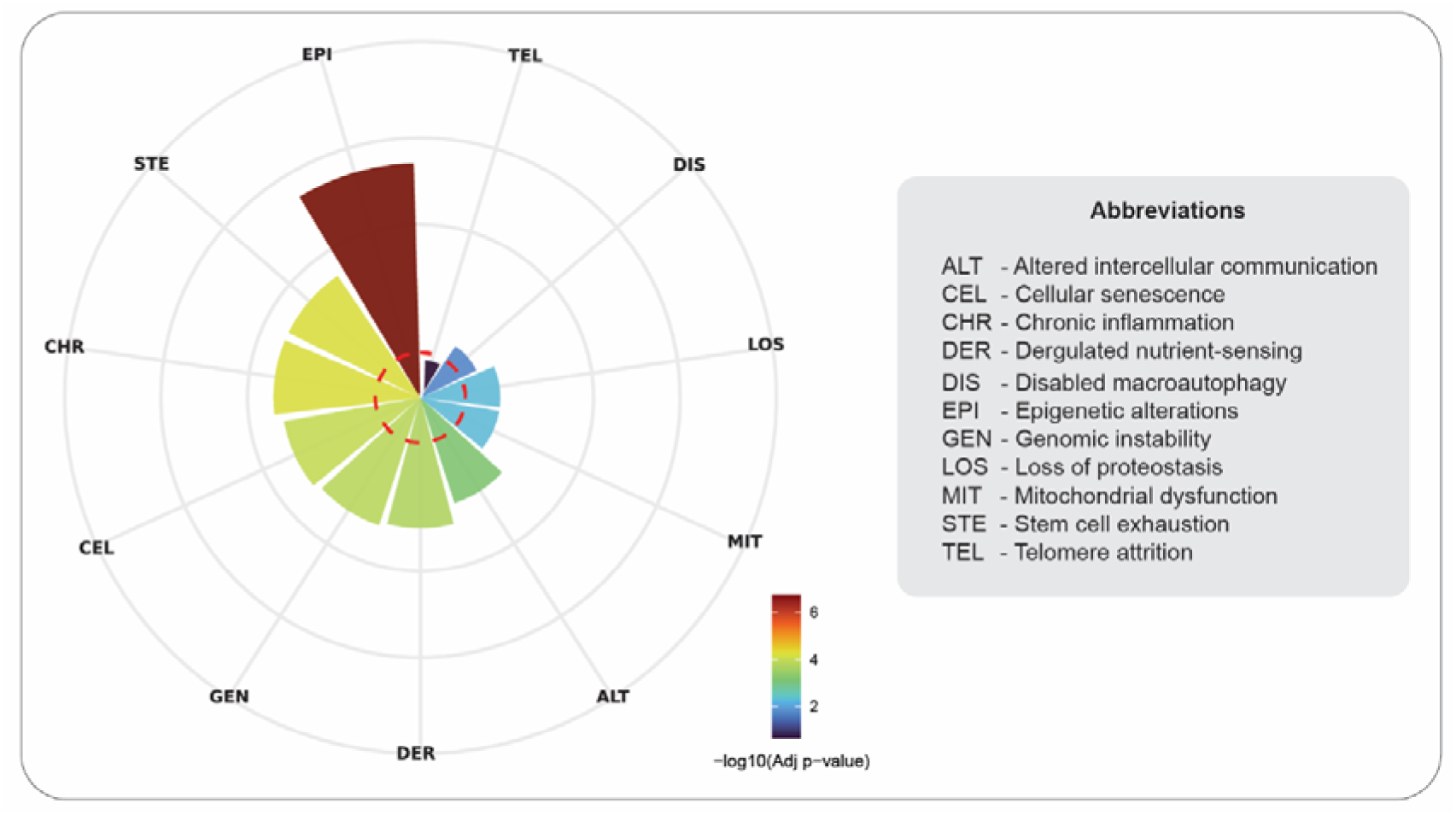
Circular plot for hallmark enrichment results for PM2.5-associated gene set using hallmark gene sets filtered for skin-specific gene expression.

## Discussion

Aging is a complex process characterized by interconnected hallmarks, that describe its progression and functional manifestation. In this study, we constructed AgingHallmarksDB, an integrative resource comprising 11 of 12 hallmarks of aging and their associated genes to facilitate a systems-level understanding of aging. AgingHallmarksDB represents one of the largest curated resources linking 3111 genes to hallmarks of aging, through a functional annotation framework capturing the biological basis of each individual hallmark and their interconnectedness at the systems level. To holistically describe aging, the resource compiles several annotations like cell type class, tissue type, exosomal presence, and regulatory interactions. These annotations enable understanding of aging mechanisms at multiple biological levels.

AgingHallmarksDB offers gene sets with varying levels of stringency for hallmark-gene associations, along with enrichment analysis for various tissue and cell type classes. The enrichment tool is powered by two complementary modules. The hypergeometric test-based ORA (Over representation analysis) enables users to assess the relevance of their gene set for aging and aging hallmarks. The gene set enrichment analysis (GSEA) framework enables the analysis of ranked gene expression data, such as log fold-change values, providing complementary insights into the association of transcriptional changes with aging and its hallmarks. In addition, the resource also supports PPI-based topology analysis and regulatory interaction analysis for transcription factor-target and kinase-substrate interactions, empowering the understanding of key nodes and regulators associated with aging and its hallmarks.

Beyond serving as an integrative repository, the hallmark-gene associations present in AgingHallmarksDB describe the hallmarks and their interconnectedness at the gene level, The genes associated with each hallmark in the resource are distinct, yet interconnected at the functional level [2,6]. This was further demonstrated through network separation and proximity analyses for aging hallmark modules, revealing an overlap or interconnectedness between hallmarks at the network level, supporting that the hallmarks are not distinct functional units but a highly interconnected biological process. Notably, the ‘Epigenetic alterations’ hallmark exhibited significant overlap with ‘Genomic instability’, ‘Cellular senescence’, and ‘Stem cell exhaustion’ in both network separation and network proximity analyses. This extensive connectivity highlights the central role of epigenetic regulation in aging, suggesting that epigenetic perturbations may influence multiple aging processes through their effects on genome stability, cellular function, and stem cell maintenance and regeneration [33]. In contrast, hallmark of ‘Telomere attrition’ had high network interconnectedness with only the hallmarks of ‘Cellular senescence’, and ‘Genomic instability’; this shows its biological significance whereby telomere-associated perturbation creates genomic perturbation that can lead to cellular senescence [34]. This high network overlap or interconnectedness among individual hallmark modules point towards existence of a larger longevity module encapsulating shared molecular mechanisms underlying aging process [35]. The common genes associated with multiple hallmarks in the curated gene set also pointed towards an interconnectedness of the hallmarks and regulation of the process. UBC, which was observed as one of the top nodes after network topology analysis of common hallmark genes, has strong association with aging [36,37]. Furthermore, regulatory interaction analysis identified several key regulators among the common hallmark genes. TP53, one of the central regulatory genes, plays a pivotal role in maintaining genomic integrity by coordinating DNA damage response and repair pathways [38]. AKT1, a serine/threonine kinase, is a major regulator of insulin signalling, nutrient sensing, and cellular metabolism, all of which are fundamental processes underlying aging [39]. In addition, multiple members of the MAPK family (MAPK1, MAPK3, and MAPK8) were identified among the common hallmark genes shared across multiple aging hallmarks, highlighting the widespread involvement of MAPK signalling in aging. This observation is consistent with a previous study demonstrating that MAPKs regulate the expression and activity of numerous proteins involved in aging, including pro-inflammatory mediators and components of the p21/p53 and p16/RB signalling pathways, which constitute the principal regulatory axes of cellular senescence [40]. Overall, these interpretations indicate larger systemic connections and core mechanistic functioning of the hallmarks in the process of aging.

The genetic interconnectedness of hallmarks, enrichment analysis, protein-protein and regulatory interactions underscore the ability of AgingHallmarksDB to uncover key molecular nodes that bridge multiple aging hallmarks and contribute to the understanding of the complex biology of aging. In addition to providing system- level analysis and facilitating its interpretation, the utility of the resource was further demonstrated through a series of practical case studies involving aging-, developmental and congenital-associated diseases and their association with the hallmarks. The analysis showed markedly stronger association of aging-associated diseases with the hallmarks in comparison to developmental and congenital diseases, which are primarily manifested from birth rather than emerging during the aging process. Furthermore, analyses of aging-associated diseases highlighted the biological relevance of the hallmark framework. For example ORA-based enrichment analysis of Alzheimer’s disease clearly established a biological association with the hallmark of ‘Loss of proteostasis’, consistent with its role as a major contributing factor in its pathogenesis [41], this was followed by ‘Deregulated nutrient-sensing’, which has been strongly associated with Alzheimer’s disease through insulin signalling and related metabolic mechanisms [42]. Similarly, network proximity analysis also supported the potential association of Alzheimer’s with ‘Chronic inflammation’, further supporting the well-established relationship between inflammatory processes and Alzheimer’s disease. Likewise, atherosclerosis exhibited a close association with ‘Chronic inflammation’ [43], consistent with the concept of inflammaging and its contribution to age-related disease development [44].

Analysis of PM2.5-induced skin aging using the GSEA framework revealed significant associations with the hallmarks of ‘Epigenetic alterations’ and ‘Chronic inflammation’. Epigenetic alterations have been closely linked to skin aging [45], and a previous study has shown that PM2.5 exposure can induce reactive oxygen species (ROS)-mediated epigenetic modifications in skin keratinocytes, contributing to skin aging and dysfunction [46]. Similarly, chronic inflammation is a well-recognized feature of pollutant-induced skin aging, as the skin serves as the primary barrier exposed to environmental PM2.5, leading to inflammatory responses that contribute to age-related skin damage [47]. These findings demonstrate the utility of AgingHallmarksDB in linking transcriptomic changes, and genetic associations with biologically relevant aging hallmarks and mechanisms.

## Conclusion

AgingHallmarksDB is an openly accessible resource that provides a systems-level framework for studying aging through the lens of aging hallmarks. It integrates aging-associated genes from seven established resources, incorporates several biological annotations, and supports hallmark enrichment analysis and network-based visualizations of aging associated genes through protein-protein and regulatory interactions. Network separation- and proximity-based analysis of hallmark associated modules showed interconnectedness through the negative separation and proximity scores. Furthermore, the utility of the resource was demonstrated through analysis of common hallmark genes and their topological importance followed by regulatory capabilities, ORA-based analysis of chronic age-related and developmental and congenital diseases, and GSEA-based analysis of PM2.5-associated skin aging. The hallmark enrichment of age-related and developmental and congenital diseases was further supported through network proximity analysis. These analyses showed that hallmarks are good indicators to understand the association of diseases and exposures with the process of aging.

AgingHallmarksDB has some notable limitations; in particular, the aging hallmark ‘Dysbiosis’ human-relevant functional annotations for reliable gene assignment are currently limited, and therefore, no genes were mapped to this hallmark. Furthermore, the genes associated with aging hallmarks were obtained from publicly available repositories, and thus, the accuracy of these gene sets depends on the quality and reliability of the source databases. Despite these limitations, AgingHallmarksDB introduces a highly integrated framework for exploring hallmark-level aging mechanisms, disease-aging relationships, and environmental aging signatures. By combining rigorously curated gene sets, up-to-date functional annotations, and extensive systems-level biological context into a centralized platform, AgingHallmarksDB provides a single-window access point for the systematic interpretation of complex aging mechanisms, thereby supporting aging and longevity research.

## Data and code availability

The data associated with this study is contained within the article, or provided as Supplementary Information files, which include an Excel file and a PDF containing additional tables, text and figures, or can be accessed via the Download option in the associated web platform https://cb.imsc.res.in/aginghallmarksdb/. The code to execute network proximity analysis is accessible through the associated GitHub repository: https://github.com/asamallab/AgingHallmarksDB

## CRediT author contribution statement

**Rahul Tiwari:** Conceptualization, Data Curation, Methodology, Visualization, Software, Formal Analysis, Writing**; Mridhula Balaji:** Data Curation, Methodology, Visualization, Formal Analysis, Writing**; Nikhil Chivukula:** Data Curation, Methodology, Visualization, Formal Analysis, Writing; **Priyotosh Sil:** Methodology, Formal Analysis, Writing**; Areejit Samal:** Conceptualization, Methodology, Supervision, Formal Analysis, Writing.

## Acknowledgements

Rahul Tiwari, Mridhula Balaji and Areejit Samal would like to acknowledge Deepak Kumar Saini and his lab members at IISc Bengaluru, for valuable discussions and suggestions. Areejit Samal would like to acknowledge funding from the Department of Atomic Energy (DAE), Government of India via Apex project to The Institute of Mathematical Sciences (IMSc), Chennai. The funders have no role in study design, data collection, data analysis, manuscript preparation or decision to publish.

