## Supplementary material for "An integrated resource for systems-level analysis of aging hallmarks and associated genes": Figure S

### Supplementary Methods

#### 1. Mapping of associated functional terms and gene sets to aging hallmarks

The twelve hallmarks of aging defined by López-Otín *et al.* [1] represent broad conceptual frameworks that encompass multiple biological processes, pathways, and molecular mechanisms underlying aging-related processes. As a result, each hallmark represents an umbrella term with diverse, context-dependent, and often overlapping interpretations, rather than discrete or directly measurable entities. Accordingly, mapping of genes to individual hallmarks cannot rely on simple one-to-one associations. Instead, it requires a systematic and integrative strategy that leverages functional annotations, pathway information, and multi-level biological evidence to accurately associate genes with specific hallmark functions. Here, we describe the systematic strategy employed to establish gene associations across the hallmarks of aging.

To enable systematic mapping of genes to the aging hallmarks, the functional annotations associated with the hallmarks were first systematically curated from different sources. The functional terms associated with each hallmark, referred to as 'Analytical Hallmarks' and provided by Open Genes [2], were used as the initial list of functional annotations. These terms offer curated functional descriptors that capture key biological processes linked to aging, and thus, provide an effective understanding of functions associated with aging hallmarks. In addition, further annotations were curated and refined based on the conceptual definitions and biological interpretations of aging hallmarks provided by López-Otín *et al.* [1, 3]. These terms were then added to the existing functional annotations for each aging hallmark, to ensure comprehensive coverage of the diverse molecular mechanisms represented within each category. Through this systematic process, 37 functional terms were associated with 11 of the 12 hallmarks of aging. The hallmark 'Dysbiosis' was not mapped to any functional term because no suitable human-relevant functional annotations were identified. The functional annotations associated with the remaining 11 aging hallmarks were then used to identify their associated genes.

##### 1.1. Systematic mapping of genes to aging hallmarks

The aging-related genes, obtained from the seven aging-related databases, namely, Aging Atlas [4], AgingReG [5], CellAge [6], CSGene [7], Digital Ageing Atlas (DAA) [8], GenAge [6], and Open Genes [2], were subsequently mapped to the aging hallmarks based on associated functional terms curated from different sources. Gene Ontology (GO)-based (https://geneontology.org/docs/download-ontology/) associations from Open Genes served as the primary reference. Here, the obtained terms were manually refined by validating existing GO terms, removing obsolete entries, and expanding the coverage with newly curated GO terms. GO terms belonging to 'Biological Process' (BP) and 'Molecular Function' (MF) categories were only retained to ensure that all assigned terms were biologically relevant within the defined functional context. For example, within the hallmark of 'Genomic instability', the associated functional term 'mitochondrial DNA instability' was mapped to GO terms such as 'mitochondrial DNA replication' (GO:0006264), along with other processes related to mitochondrial DNA synthesis and its regulation (Supplementary Table S2). This framework was further extended to pathway-level annotations by manually incorporating KEGG [9] and Reactome pathways [10], which were assigned to the same associated functional terms using a similar curation strategy based on biological relevance. In sum, this integrative framework provides a functional basis for each hallmark, resulting in a systematic and biologically meaningful gene-to-hallmark mapping.

In order to map genes to the hallmarks, each gene was mapped to the associated functional terms based on its own functional annotations from GO, KEGG, and Reactome. For GO-based mapping, the 'gene2go.csv' file was used as the primary source, restricted to human genes using the taxonomic identifier '9606', and complemented with manually incorporated GO terms from the QuickGO database (https://www.ebi.ac.uk/QuickGO/annotations) to ensure completeness. All available evidence code annotations were retained, and those corresponding to experimental annotations, including 'EXP', 'IDA', 'IPI', 'IMP', 'IGI', 'IEP', 'HTP', 'HDA', 'HMP', 'HGI', and 'HEP', along with manually curated annotations such as 'TAS', 'NAS', and 'IC', were used to identify a high confidence association subset (Supplementary Table S4). The pathway-level associations were incorporated using data from Reactome and KEGG. Reactome-based associations were derived from the 'NCBI2Reactome_All_Levels.csv' file, wherein all available entries were retained, with those supported by 'TAS' evidence and restricted to *Homo sapiens* used to identify high confidence associations. In parallel, KEGG-based gene-pathway relationships were retrieved using the KEGG API, filtered for human-specific entries, and standardized to Entrez Gene identifiers using the mygene package (https://pypi.org/project/mygene/) (Supplementary Table S3). Finally, genes associated with sirtuin pathways as documented by the HUGO Gene Nomenclature Committee (HGNC; https://www.genenames.org/) were manually assigned to the aging hallmark 'Deregulated nutrient-sensing', as these genes were not adequately captured through GO, KEGG, or Reactome mappings. Subsequently, the intersection of these mapped genes with the aging-related genes from the seven databases was used to identify the hallmark-associated genes. Collectively, this integrated framework resulted in a comprehensive hallmark-to-gene dataset, enabling a systematic and biologically meaningful assignment of genes to their respective aging hallmarks. Furthermore, this exercise provided a clearer understanding of the underlying functional and biological mechanisms associated with aging hallmarks.

**2. Additional characterization of hallmark-associated genes**

To obtain a systems-level understanding of hallmark-associated genes, multi-layer biological annotations were integrated, including single-cell expression, exosomal presence, and regulatory interactions such as transcription factor (TF)-target and kinase-substrate relationships. This enabled a more comprehensive interpretation of gene function and context across aging hallmarks. Single-cell expression data was obtained from the 'rna_single_cell_type.tsv' file provided by the Human Protein Atlas (https://www.proteinatlas.org/) [11]. The Ensembl gene identifiers were standardized to Entrez identifiers using biomaRt [12], and unmapped or ambiguous entries were excluded. Thereafter, biologically meaningful gene-cell type associations were identified based on expression cut-off of nCPM > 1 (https://www.proteinatlas.org/humanproteome/single+cell/single+cell+type/method). The 'rna_single_cell_type_cell_types.tsv' and 'rna_single_cell_clusters.tsv' files were further utilized to characterize the associated cell types into broader cell type class and tissues, respectively.

Exosome-associated RNA datasets were obtained from 'circRNAs_anno.csv' and 'longRNAs_anno.csv' files from exoRBase (v3.0) [13], and gene symbols were standardized to Entrez identifiers using mygene package. Subsequently, the hallmark genes found in this dataset were annotated as present in exosomes. The TF-target interactions were sourced from DoRothEA dataset available on OmniPath [14], and filtered to obtain interactions with high confidence scores of A to C. Lastly, the kinase-substrate phosphorylation events were curated from the EnzSub dataset available in OmniPath [14].

**3. Network separation and proximity analysis**

Network separation and proximity analysis describes human interactome level interactions between hallmark-hallmark pairs or hallmark-disease pair, giving a descriptive systemic insight of the process. Here network separation is used to analyse the separation between the hallmark pairs on the human interactome, on the other hand proximity is employed to firstly understand the proximity between the hallmark pairs followed by proximity between the hallmark and disease pairs (age and developmental and congenital disease). The computations were performed using the Python package NetMedPy (version 0.1.171) by leveraging a comprehensive manually curated human protein-protein interaction (PPI) network [15].

#### 3.1. Compilation of human protein-protein interaction network

A large-scale human PPI network was assembled to support the computation of network proximity, integrating multiple high-quality interactomes to achieve broad coverage. The first dataset included the PPI network reported by Gan *et al.* [16], which comprised 18505 proteins as nodes and 327924 interactions as edges. This dataset was complemented by the HI-union interactome from the Human Interactome Atlas (https://interactome-atlas.org/) [17]. As this dataset was provided in Ensembl gene identifiers, conversion to Entrez identifiers was performed using mygene (https://pypi.org/project/mygene/). Following the removal of duplicate interactions and self-loops, the HI-union dataset contained 9053 nodes and 63669 unique interactions. Additionally, the human PPI network curated by Ruiz *et al.* [18], which integrates interaction data from multiple sources, was included, comprising 17660 nodes and 387626 edges. The three networks or datasets were subsequently combined by taking the union of nodes and interactions, with duplicate edges removed to ensure consistency. The resulting integrated interactome comprised 19572 unique proteins and 541092 high-confidence interactions. It was observed that the largest connected component (LCC) of this constructed PPI network comprised 19545 nodes and 541065 edges, and the LCC was used as the background network for all proximity computations.

#### 3.2 Statistical significance of hallmark-associated LCCs

The statistical significance of LCC for each hallmark-associated gene set was assessed using NetMedPy. The observed LCC size was compared with a null distribution generated from 1000 degree-preserving randomization (log-binning). Statistical significance was quantified by a z-score, which for observed LCC size n is given by:

$$z = \frac{n - \mu_n}{\sigma_n} \tag{1}$$

where $\mu_n$ is the expected LCC size chosen randomly, and $\sigma_n$ is the LCC standard deviation across randomized ensemble. Higher positive values indicates that the hallmark-associated genes formed a significantly more connected module within the human interactome than expected by chance.

#### 3.2 Network separation and its implementation

Network separation analysis was conducted using NetMedPy python package. All computations were restricted to LCC containing 19545 nodes and 541065. To understand the overlap or interconnectedness between a pair of aging hallmarks modules A and B its separation $S_{AB}$ could be calculated by

$$S_{AB} = d_{AB} - \frac{d_{AA}+d_{BB}}{2} \tag{2}$$

where $d_{AB}$ represents the average of the shortest path length between all genes of hallmarks A and B. Thus, $S_{AB}$ compares the shortest distance between hallmark-associated modules A and B with their respective within-module distances $d_{AA}$ and $d_{BB}$. A negative separation value indicates that the two modules are more closely connected to each other than expected from their internal connectivity, reflecting topological overlap within the human interactome.

#### 3.3 Network proximity and its implementation

Network proximity analysis was conducted using NetMedPy [15]. All computations were restricted to the LCC containing 19545 nodes and 541065 edges of the integrated PPI network to ensure reliable shortest-path distance estimation. Each aging hallmark comprises a curated set of associated genes, which were aggregated to define the hallmark-specific gene set $H$, while the case study specific candidate gene set $D$ was constructed as described in the Methods section in the Main text. The network proximity was calculated using the average minimum shortest path length ($d_c$) in its asymmetric formulation [15, 19]. The closest distance between the aging hallmark gene set $H_k$ and the case study specific gene set $D$ is defined as,

$$d_c(H_k, D) = \frac{1}{|D|}\sum_{y \in D} min_{x \epsilon H_k} d(x, y) \tag{3}$$

Here, $d(x, y)$ represents the shortest-path distance between node $x$ (aging hallmark genes) and node $y$ (case study specific genes) within the PPI network. Smaller values of $d_c$ reflect a closer topological relationship between the aging hallmark gene set and the case study specific gene set $D$.

Simultaneously, to quantify the overlap between hallmark-associated gene sets $H_A$ and $H_B$ on within the PPI network, symmetric network proximity analysis was performed, as both gene sets represented aging hallmarks. This analysis first computed the closest-path distances $d_c(H_A, H_B)$ and $d_c(H_B, H_A)$ using the closest-distance metric defined in Equation (3), followed by the calculation of their symmetric proximity score.

$$d_c(H_A, H_B) = \frac{1}{|H_B|}\sum_{y \in H_B} min_{x \epsilon H_A} d(x, y) \quad (4)$$

Following to that it calculates another $d_s(H_B, H_A)$, which is given by,

$$d_s(H_A, H_B) = \frac{d_c(H_A,H_B)+d_c(H_B,H_A)}{2} \quad (5)$$

The symmetric proximity score, $d_s(H_A, H_B)$ is defined as the average of the closest-path distances from hallmarks $H_A$ and $H_B$ and from $H_B$ to $H_A$. This bidirectional measure provides an unbiased estimate of the overall network proximity between two hallmark-associated gene sets within the human protein-protein interaction network, where lower values indicate greater network proximity.

#### 3.3. Statistical significance assessment

To determine whether the observed network proximity between the aging hallmark gene set $H_k$ and the case study specific gene set $D$ was lower than expected by chance, a degree-aware null model was applied using NetMedPy [15]. Randomization was carried out using a logarithmic binning approach, maintaining the size of the original gene set while approximating its degree distribution. For each aging hallmark, 1000 randomized gene sets $H_k^{(r)}$ were generated, and their network proximity to the case study specific gene set $D$ was computed using the same asymmetric average shortest path length ($d_c$) metric described above. This procedure produced an empirical null distribution of proximity values. The statistical significance of the observed proximity $d_c(H_k, D)$ was then assessed by calculating a z-score:

$$z = \frac{d_c(H_k,D)-\mu_{rand}}{\sigma_{rand}} \quad (6)$$

Here, $\mu_{rand}$ and $\sigma_{rand}$ represent the mean and standard deviation values obtained from the randomized samples, respectively. $\mu$ and $\sigma$ for symmetric condition is calculated using a null model that considers definition of $d_s$ in equation (5).

A one-tailed $p$-value was then calculated to determine whether the observed proximity was significantly lower than expected under the null model. This $p$-value corresponds to the proportion of randomized proximity values that are less than or equal to the observed value. Negative z-scores in combination with $p < 0.05$ were interpreted as evidence of statistically significant proximity between the different hallmark gene sets and the case study specific gene sets.

### Supplementary Figures

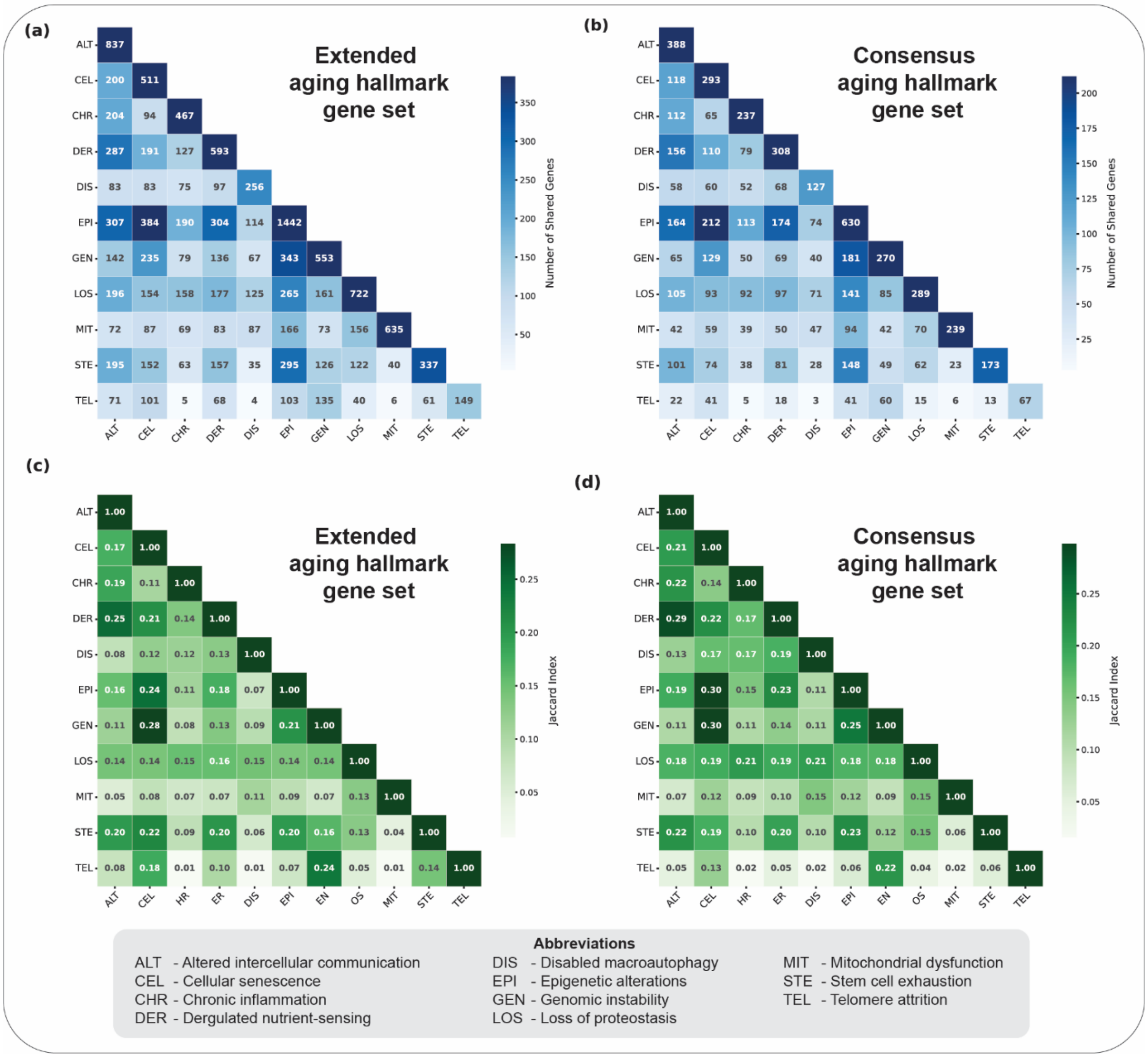


**Figure S1.** Pairwise analysis of gene overlaps between different aging hallmarks. **(a)** Pairwise overlap between extended aging hallmark-associated gene sets. **(b)** Pairwise overlap between consensus aging hallmark-associated gene sets. **(c)** Pairwise Jaccard index between extended aging hallmark-associated gene sets. **(d)** Pairwise Jaccard index between consensus aging hallmark-associated gene sets.

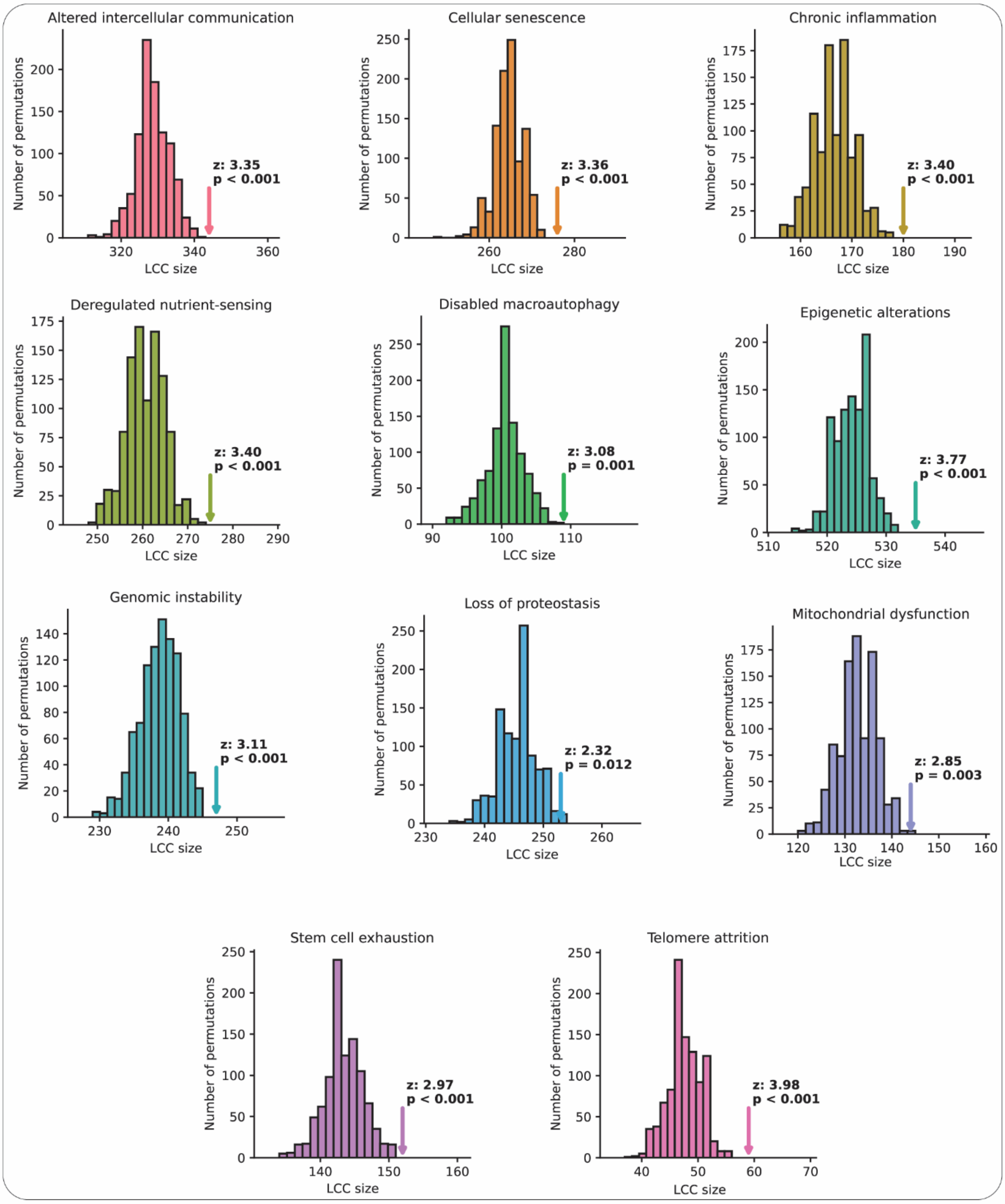


**Figure S2.** The size of LCC obtained for each hallmark and its associated significance, compared with distribution of LCC formed by randomized control groups. Every LCC represents a highly connected 'module' for corresponding hallmark of aging.
